# Global priorities for conservation of reptilian phylogenetic diversity in the face of human impacts

**DOI:** 10.1101/723742

**Authors:** Rikki Gumbs, Claudia L. Gray, Monika Böhm, Michael Hoffmann, Richard Grenyer, Walter Jetz, Shai Meiri, Uri Roll, Nisha R. Owen, James Rosindell

**Affiliations:** Department of Life Sciences, Imperial College London, Silwood Park Campus, Ascot, Berkshire, SL5 7PY, United Kingdom; Science and Solutions for a Changing Planet DTP, Grantham Institute, Imperial College London, South Kensington, London; EDGE of Existence Programme, Zoological Society of London, Regent’s Park, London, United Kingdom; Institute of Zoology, Zoological Society of London, Regent’s Park, London, United Kingdom; Conservation and Policy, Zoological Society of London, Regent’s Park, London, United Kingdom; School of Geography and the Environment, University of Oxford, Oxford, OX1 3QY, United Kingdom; Ecology and Evolutionary Biology Department, Yale University, 165 Prospect Street, New Haven, CT 06511, USA; School of Zoology, Tel Aviv University, 6997801, Tel Aviv, Israel; Steinhardt Museum of Natural History, Tel Aviv University, 6997801, Tel Aviv, Israel; Mitrani Department of Desert Ecology, Ben-Gurion University of the Negev, Midreshet Ben-Gurion 8499000, Israel; On The EDGE Conservation, London, United Kingdom

## Abstract

Phylogenetic Diversity (PD) is increasingly recognised as an important measure that can provide information on evolutionary and functional aspects of biodiversity for conservation planning that are not readily captured by species diversity. Here we develop and analyse two new metrics that combine the effects of PD and human encroachment on species range size — one metric valuing regions and another enabling species prioritisation. We evaluate these metrics for reptiles, which have been largely neglected in previous studies, and contrast these results with equivalent calculations for all terrestrial vertebrate groups. We find that high human impacted areas unfortunately coincide with the most valuable areas of reptilian diversity, more than expected by chance. We also find that, under our species-level metric, the highest priority reptile species score far above the top mammal and bird species, and they include a disproportionate number of species with insufficient information on potential threats. Such Data Deficient species are, in terms of our metric, comparable to Critically Endangered species and may require urgent conservation attention.

## Introduction

We are in the midst of a global biodiversity crisis^1, 2^ with severely limited resources for conservation action^3^. At current extinction rates, we are set to experience unprecedented losses of species and their Phylogenetic Diversity (PD). PD is the sum of the phylogenetic branch lengths connecting a set of species to each other across their phylogenetic tree, and measures their collective contribution to the tree of life^4, 5^. PD is increasingly recognised as an important component of global biodiversity^6, 7^ with value for human well-being^4, 8, 9^. As PD extends beyond the simple counting of species to quantify the amount of variation across a set of species^4^, it is a valuable tool for differentiating among species and regions for conservation prioritisation^5, 10–12^.

A large body of literature has explored how PD can be conserved across mammals and birds^5, 7, 10, 13–16^ but reptiles remain poorly studied in global conservation schemes^17^, despite comprising ^~^30% of terrestrial vertebrate species richness^18^. Almost one in five reptile species are threatened with extinction^19^ and reptile populations have suffered average global declines of around 55% between 1970 and 2012^20^. Existing protected areas and global conservation schemes represent reptiles poorly compared with birds and mammals^21^. Consequently, there is a pressing need to assess all reptiles to enable targeted conservation and allow the incorporation of reptiles into global analyses of conservation priorities.

There are several methods available for mapping imperilled PD ^7, 12, 13, 22, 23^ and, in lieu of explicit extinction risk data, small range size has often been used to identify regions of high conservation value^12, 13^. However, whilst these methods prioritise highly irreplaceable regions, they do not incorporate spatial measures of vulnerability, such as human impact, thus limiting their potential practical application in conservation planning^24, 25^. Unfortunately, while range data to roughly 100 km scale are now available for 99% of reptiles^21^, up-to-date extinction risk data (i.e. published in the past ten years^26, 27^) are available for less than half of reptile species^27^. In the absence of comprehensive extinction risk assessments for all reptiles, range data must be combined with existing environmental data to determine spatial vulnerability^28–30^.

The Human Footprint index (HF)^30, 31^ is the most comprehensive and high-resolution dataset of human pressures on global environments. It combines eight variables—including crop and pasture land, extent of built environments, human population density and night-time lights–which measure direct and indirect impacts of humans on the environment^30^. Such comprehensive global maps of cumulative human pressures have been shown to be better predictors of species distributions than biological traits^32^ and are a strong predictor of species extinction risk^33^. However, to our knowledge, no measure of human impact—such as the Human Footprint—has previously been explicitly incorporated into methods to value and prioritise the conservation of global vertebrate PD. Here, we present two new metrics combining human encroachment (to measure vulnerability), and range size (to measure irreplaceability), to identify high value regions and high priority species for conserving reptile PD. For comparison, we also calculate these metrics at the global scale for all tetrapod clades.

## Methods

### Data

We used updated reptile distribution polygons from the Global Assessment of Reptile Distributions (GARD)^21^. We used published phylogenies for lepidosaurs (lizards, snakes and the tuatara)^34^, crocodilians^35^ and turtles^36^. The crocodilian and turtle phylogenies used were single, consensus, fully-resolved phylogenies. To capture phylogenetic uncertainty around the taxonomically imputed lepidosaur phylogenies, we randomly sampled 100 fully-resolved phylogenies from a distribution of 10,000 trees^34^ and used each phylogeny in our analyses to generate median values of PD and PD-based metrics for each grid cell using a Mollweide equal area projection at 96.5 x 96.5 km grid cell resolution^21^. We matched the species in each phylogeny to the distribution data using the taxonomy from the July 2018 version of the Reptile Database^18^. For our spatial analyses we included only species with both phylogenetic and distribution data (9,862 species or 91% of total reptilian diversity; Supplementary Table 1).

We extracted a random sample of 100 phylogenetic trees from published phylogenies for amphibians^37^, birds^13^ and mammals^38^ and spatial data, as polygon shapefiles, for amphibians and mammals from IUCN^27^ and for birds from BirdLife International^39^. These distribution data were subset to contain only native and resident or breeding ranges. As with reptiles, for our spatial analyses we included only species with both phylogenetic and distribution data (5,786 amphibians (75.5% of species); 9,274 birds (84.5%); 4,386 mammals (77%) - ^~^84% of all tetrapods, including reptiles; Supplementary Table 1) and calculated median values of PD and PD-based metrics for each grid cell.

We used the 2009 Human Footprint index (HF)^30^—the most up-to-date HF dataset—to designate spatial patterns of human pressure. The HF index evaluates each grid cell based on the intensity of eight measures of human pressure (built environments, crop land, pasture land, human population density, night-time lights, railways, roads, navigable waterways), weighted according to estimates of their relative levels of human pressure^30, 31^, and assigns an HF value between 0 (lowest human pressure) and 50 (greatest human pressure) to each cell^30^. We resampled the HF data from its original 1 x 1 km resolution to our 96.5 x 96.5 km grid.

### Spatial value metric for conserving PD

As small range size is linked to elevated extinction risk^29, 40^, if small-ranged reptiles are clumped together on the tree of life, with no shared branches also subtended by a wide-ranging species, a disproportionately large amount of PD may be at risk of extinction. To examine whether small range size is phylogenetically conserved in this manner, we calculated Pagel’s lambda^41^ for crocodilians, turtles, and lepidosaurs separately and—within lepidosaurs—for lizards, amphisbaenians, and the tuatara (hereafter collectively ‘lizards’) and for snakes independently, to remove the biased caused by large range sizes of snakes from the analysis of lizard distributions^21^. Pagel’s lambda provides an estimate of how phylogenetically conserved a trait is across a phylogeny, with scores close to 1 indicating a trait is extremely clumped on the phylogeny, whereas scores close to 0 indicate a trait to be randomly dispersed throughout the phylogeny^41^.

To map global patterns of reptilian PD, for each grid cell occupied by at least one species, we summed the lengths of all branches between root and tips for each species in the grid cell. As the branch lengths are time-calibrated, the resulting values represent the PD, as units of time, present in each grid cell. To account for the internal branches connecting crocodilians, turtles and lepidosaurs when mapping PD for all reptiles, we used published divergence estimates between each clade pair^42^.

We summed the branch lengths of the turtle and crocodilian phylogenies, and combined these with the additional inferred PD and the median summed branch lengths from the 100 lepidosaur phylogenies to estimate total global reptilian PD. Though crocodilians were included in analyses of all reptiles, we do not report their individual results because they comprise of only 25 species^18^.

We explored the relationship between PD and richness for each reptile group using Pearson’s correlation corrected for spatial autocorrelation in the R package ‘Spatialpack’^43, 44^, with conservative Bonferroni correction for multiple testing. To identify global variation in the relationship between PD and richness, we calculated the residuals from a linear regression of richness against PD for all grid cells. We consider grid cells harbouring more PD than expected for the observed richness to represent regions of disproportionately phylogenetically diverse species compositions.

For later comparison with our own PD-based spatial metric, we calculated three additional metrics: the species-based metric Weighted Endemism (WE), which provides a measure of range-size-weighted species richness^12, 21^, and two PD-based extensions of Weighted Endemism: Evolutionary Distinctness Rarity (EDR)^13^ and Phylogenetic Endemism (PE)^12^ (Supplementary Table 2).

A key difference between the two PD-based metrics, Evolutionary Distinctness Rarity and Phylogenetic Endemism, is in their treatment of species ranges: Evolutionary Distinctness Rarity treats all species ranges as spatially independent whereas Phylogenetic Endemism accounts for the spatial overlap of species. We suggest that Evolutionary Distinctness Rarity and Phylogenetic Endemism therefore better represent the potential loss due to differing drivers. Evolutionary Distinctness Rarity represents the amount of Evolutionary Distinctiveness imperilled by *species*-specific threats (e.g. targeted hunting); the losses are species focused because only range size (and not range overlap with other species) is accounted for. In contrast, Phylogenetic Endemism represents the amount of phylogenetic diversity attributed to a particular unit of *space*, reflecting the impact of landscape-level threats (e.g. habitat loss); having additional descendent species in the same size region makes no difference to extinction risk of phylogenetic branches because loss of the region would impact all those species together. As most threats to tetrapod species are present at the landscape-level (e.g. agriculture, logging and livestock production)^45–47^, we hereafter report and develop analyses based on the Phylogenetic Endemism metric.

To assess the overlap between regions of high Phylogenetic Endemism and high human pressure, we identified the grid cells in the top 10% of all grid cells for reptilian Phylogenetic Endemism (hereafter “high value grid cells”) and calculated the proportion of the high value grid cells that are also deemed to be under ‘high’ or ‘very high’ human pressure (Human Footprint ≥ 6)^30^. As Human Footprint value of 4 equates to the human pressure of pasture lands^48, 49^, ours is a conservative estimate of intense human pressure^30^. We randomised the distribution of grid cells under high or very high human pressure across all terrestrial cells and recalculated the proportion of high value grid cells now considered to be under high or very high human pressure. We repeated this randomisation 1,000 times to generate a distribution of randomised scores for comparison with the observed proportion of overlap.

Whilst Phylogenetic Endemism incorporates the intrinsic threat of small range size into the calculation of grid cells for conservation of unique evolutionary history, it does not measure the myriad extrinsic threats present. We therefore incorporated the Human Footprint (HF) index^30, 31^ as a measure of vulnerability.

To calculate an adjusted range size value for each species in relation to HF we first linearly scored each terrestrial grid cell between 0 and 1 according to which of the five approximately equally distributed classes of HF it belonged: HF-adjusted range size of 1 = ‘no pressure’ (HF = 0), the entire grid cell is retained; 0.8 =‘low pressure’ (HF = 1-2); 0.6 = ‘moderate pressure’ (HF = 3-5); 0.4 = ‘high pressure’ (HF = 6-11); 0.2 = ‘very high pressure’ (HF = 12-50) ^30^. A ‘very high pressure’ grid cell is therefore equivalent to 0.2 of a complete grid cell, to reflect the high human pressure and therefore likely greatly reduced remaining suitable habitat within that cell for species to persist. Though the true proportion of remaining suitable habitat will differ across grid cells of equal Human Footprint and will also be species-specific, our scoring of grid cells based on Human Footprint provides a relative scale representing human pressure under the assumption that increased human pressure equates to less remaining suitable habitat. The new “HF-adjusted range size” of a species is given by the sum of HF-adjusted grid cell size for all cells across which a species is distributed. It can be thought of as an effective range size, which will be much smaller than the true range if large parts of it coincide with high levels of human pressure. Previous analyses have used fine-scale environmental data to estimate range loss across species under scenarios of change^50^, and combined these with phylogenetic data on a regional scale for a relatively small clade^51^. However, such fine-scale habitat association and environmental requirement data are lacking for the majority of reptiles and preclude such an analysis at this time.

We used these HF-adjusted range sizes to calculate a new spatial PD metric, derived from PE, which we term Human Impacted Phylogenetic Endemism (HIPE). This approach apportions the PD of each branch of the phylogeny according to each grid cell’s contribution to the total adjusted range of the species (Supplementary Table 2). When a branch is found either in one grid cell or in multiple grid cells of the same HF-adjusted grid cell size, HIPE is equivalent to Phylogenetic Endemism in apportioning PD. However, when a branch occurs in grid cells of variable human impacts, PD is apportioned by the relative contribution of the Human Footprint-adjusted grid cells, so that those with lower human impact (higher HF-adjusted grid cell size) receive a greater proportion of PD to reflect their higher present value. Consequently, branches which are entirely distributed across grid cells of high human impact contribute a greater proportion of PD to highly impacted grid cells than branches which also occur in grid cells under low human impact.

Consider a grid cell under high human impact (HF-adjusted range size = 0.2) where only two branches are present, both comprising 10 MY of PD. Both branches also occur in one other grid cell, branch A in a low impact grid cell with a HF-adjusted range size of 1 (for a total HF-adjusted range of 1.2 grid cells) and branch B in a high impact grid cell with a HF-adjusted range size of 0.2 (total HF-adjusted range of 0.4 grid cells). Under traditional Phylogenetic Endemism, the grid cell receives 50% of the PD from each branch (5 MY) as it comprises 50% of the total distribution of the branch (one of two grid cells). Under HIPE, however, branch A would apportion only 1/6th (1.667 MY) of its PD to the grid cell as it comprises only 1/6th of the total HF-adjusted range (0.2 of total 1.2 range), with the remaining 5/6th of the PD being apportioned to the grid cell with a HF-adjusted range size of 1. Conversely, as branch B occurs only in two grid cells of HF-adjusted range size 0.2, the grid cell comprises 50% of the HF-adjusted range of the species (0.2 of total 0.4 range) and is apportioned 50% of the PD of the branch (5 MY) (Supplementary Table 2; Supplementary Figure 1).

HIPE increases the relative importance of grid cells under low human impact as well capturing cells with high endemic PD. It is therefore important for conservation planning to highlight which of the high value regions (based on HIPE) are driven by endemic PD in areas of high vs. low human impact, as the two extremes are likely to require different conservation action. We partitioned global patterns of HIPE by human impact, highlighting regions of high HIPE and high human impact (HF ≥ 6) and regions of high HIPE and low human impact (HF < 3).

We mapped HIPE for all reptile groups individually and for all reptiles combined. To determine the regions where reptiles provide the greatest contributions to global patterns of tetrapod HIPE, we also calculated HIPE for mammals, birds, amphibians and for tetrapods as a whole. We then calculated the proportions of observed HIPE for all tetrapods that were contributed by each tetrapod clade. We present HIPE scores in MY/km^2^, where the adjusted range size represents the area across which the scores are divided (e.g. a 96.5 x 96.5 km grid cell with a HF-adjusted grid cell size of 0.2 is considered to comprise 1/5^th^ of the area of an entire grid cell).

We ran spatially-corrected correlations between HIPE, Phylogenetic Endemism and Evolutionary Distinctness Rarity to test the extent to which these measures capture the same global patterns. We also ran a spatially-corrected correlations test for relationships between global HIPE patterns among reptile groups and between reptiles and other tetrapods, all with Bonferroni correction for multiple testing.

### Species prioritisation metric for conserving PD

We estimated the total PD of reptiles by summing the branch lengths of the crocodilian and turtle phylogenies and adding these to the summed branch lengths for each of the 100 lepidosaur phylogenies to generate a distribution of 100 total reptilian PD values. We compared this distribution with that for other tetrapod classes, which we generated by summing the branch lengths of the 100 random phylogenies for amphibians, birds and mammals. We compared the distributions of PD scores using ANOVA and applied Tukey’s HSD test to identify pairwise differences between tetrapod classes. The branch lengths were summed for all phylogenies prior to the removal of species with no spatial data to limit the impact of differing availability of spatial data across the different classes.

To identify species that should be prioritised to preserve unique evolutionary history, we devised a new metric built around the main component common to both PE and EDR: terminal branch length (TBL). Terminal branches are those which connect the species (tips) to the internal branches of the phylogeny and are the only component of a phylogeny unique to each species. The length of these terminal branches represents the divergence time (in millions of years for time-calibrated phylogenies) between a species and their closest relatives.

We defined Terminal Endemism (TE) as the terminal branch length of a species multiplied by 1/the number of grid cells occupied by the species. If a species is found in only one grid cell then its loss from that grid cell would result in the loss of its entire terminal branch. The TE of a species is implicitly calculated when calculating both Evolutionary Distinctness Rarity and Phylogenetic Endemism and represents the unique contribution of the species to the total for each metric. We posit that, as a species focused measure, TE circumvents the differences between Evolutionary Distinctness Rarity and Phylogenetic Endemism and retains the most essential component of each.

To incorporate HF, we developed a counterpart to TE, ‘Human Impacted Terminal Endemism’ (HITE). This metric is given by the terminal branch length of a species divided by its Human Footprint-adjusted range size (see above). For example, a species with a terminal branch length of 10 MY that is found in two grid cells, with HF-adjusted grid cell sizes of 0.2 and 1 would receive a HITE score of 10*(1/(1+0.2)) = 8.34. Under standard Terminal Endemism the same species would receive a lower score of 5: (10*(1/2)). HITE therefore increases in response to terminal branches occurring in grid cells under high human impact.

We calculated the terminal branch lengths, HF-adjusted range size and HITE for all tetrapods and ranked the species from each clade to identify the species with the highest HITE scores. We highlight tetrapod species which are either unassessed or listed as Data Deficient by the IUCN, but have a high HITE score. These are species that, due to their high irreplaceability and extremely restricted and human-impacted range, are priorities for conservation assessment. Finally, we compared HITE scores for tetrapods across IUCN Red List categories, using ANOVA and Tukey’s HSD test, to determine the relationship between HITE scores, data deficiency, and extinction risk across reptiles and all tetrapods.

To estimate how much reptilian PD may be lost if all threatened species were to become extinct, we dropped all species listed in threatened categories on the IUCN Red List (i.e. Vulnerable, Endangered and Critically Endangered) from their respective phylogenies and calculated the reduction in total PD. For lepidosaurs we did this for all 100 phylogenies to generate a distribution of values. To determine whether this potential loss of PD was greater than if extinction risk was randomly distributed across the reptilian tree of life, we then selected 100 random sets of species corresponding to an equal number of species as those observed to be threatened and dropped them from their respective phylogenies. We then compared the distribution of potential PD loss from species observed to be threatened with the distribution generated from randomised extinction using a paired t-test.

As it is likely that a significant proportion of unassessed and Data Deficient species are also threatened with extinction^52, 53^, these estimates of loss of PD are conservative. To explore how data deficiency affects potential losses of PD across data-poor regions of the tree of life, we selected a poorly-known squamate genus as a case study. We estimated the amount of PD lost under different scenarios of phylogenetic relationships and extinction risk for *Dibamus*, one of the least-known reptilian genera (and the sister clade to all other squamates). First, we estimated the amount of PD represented by the *Dibamus* species included in the phylogeny despite lacking genetic data across our random selection of 100 lepidosaur phylogenies. Second, we estimated how much PD would be lost under three extinction scenarios for *Dibamus*: 1) only a single unassessed or Data Deficient species becomes extinct; 2) a random number and selection of unassessed or Data Deficient species become extinct; and 3) all unassessed and Data Deficient species become extinct.

## Results

### Spatial value metric for conserving PD

Range size is weakly phylogenetically conserved across lepidosaurs (λ= 0.373, p ≪ 0.0001; Supplementary Figure 2), and in lizards and snakes independently (lizards: λ= 0.485, p ≪ 0.0001; snakes: λ= 0.345, p ≪ 0.0001). However, range size is not significantly conserved across turtles (λ= 0.12, p = 0.03) or crocodilians (λ= 0.048, p = 0.815), following Bonferroni correction for multiple testing (adjusted p-value threshold = 0.01), likely due to the low species richness of both clades.

Reptilian PD is largely concentrated throughout the tropics (Figure 1, Supplementary Figure 3), and is strongly correlated with species richness on a global scale for all reptiles (r = 0.948, e.d.f. = 23.7, p ≪ 0.0001), lizards (r = 0.920, e.d.f. = 21.1, p ≪ 0.0001), snakes (r = 0.899, e.d.f. = 24.4, p ≪ 0.0001) and turtles (r = 0.873, e.d.f. = 28.1, p ≪ 0.0001). Lizard PD is high across Southeast Asia, the Amazon basin and Australia (Figure 1a). Large concentrations of snake PD are found in Malaysia and Indonesia (Figure 1d), whereas the greatest concentrations of turtle PD are found across the Amazon Basin (Figure 1g), despite turtle richness peaking in the Ganges Delta.

**Figure 1:**
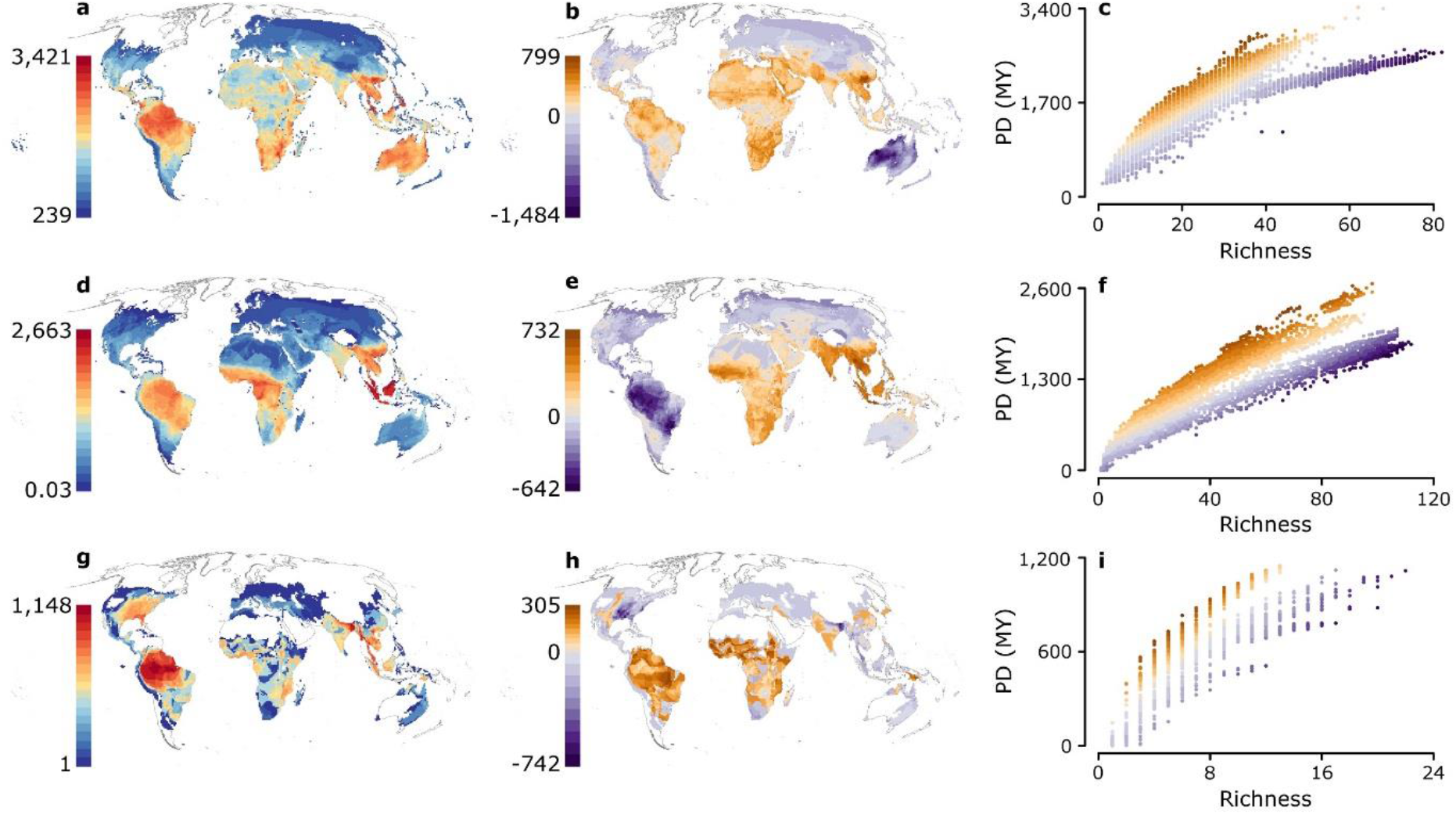
Global patterns of reptilian phylogenetic diversity (PD). Cumulative PD, in millions of years (MY) (left), Middle: residual PD per grid cell, in MY, (warm colours: more than expected given richness, cold colours: less than expected given richness), Right: the relationship between richness and PD across all grid cells for lizards (a-c), snakes (d-f), and turtles (g-i).

The greatest levels of high lizard PD, compared with species richness (residuals of PD vs. richness), are in mainland Southeast Asia, whereas regions with the lowest levels of residual PD occur across Australia, where richness is highest (Figure 1b-c). The largest accumulations of snake PD for a given richness occur in mainland Southeast Asia, and the lowest coincide with the species-rich Amazon Basin and Atlantic coast of Brazil (Figure 1e-f). The greatest accumulations of turtle PD for a given richness occur across subtropical West and Central Africa and the Amazon Basin, with lowest accumulations occurring where species richness is highest: the Ganges Delta and Eastern USA (Figure 1h-i).

Phylogenetic Endemism (PE) and Evolutionary Distinctness Rarity (EDR) for reptiles are highly correlated at the global scale (r = 0.975, e.d.f. = 537, p ≪ 0.0001) and both are highly correlated with the non-phylogenetic measure of Weighted Endemism (WE; both r > 0.93; Supplementary Figure 4).

Almost three-quarters (74%) of high-value grid cells of PE (i.e. top 10% ranking grid cells) are in regions of high or very high human pressure (Human Footprint ≥ 6), whereas just 5% of high PE grid cells coincide with regions of low or no human pressure (HF < 3). The strong association of regions of high PE with those of high human impact is surprising, considering the two are, in theory, independent. Indeed, when we randomise human pressure across grid cells, less than half (49%) of high-value grid cells coincide with high or very high human pressure, and 20% coincide with regions of low or no human pressure. High reptilian PE coincides with very high human pressure (HF > 11) across the tropics—particularly in India, Caribbean islands, the Atlantic Coast of Brazil, and Southeast Asia— and the Mediterranean coast and areas of the Middle East (bold red regions, Figure 2a). Regions of high PE and very low human pressure (HF < 1) are largely restricted to the Amazon Basin and central Australia (bold blue regions, Figure 2a).

**Figure 2:**
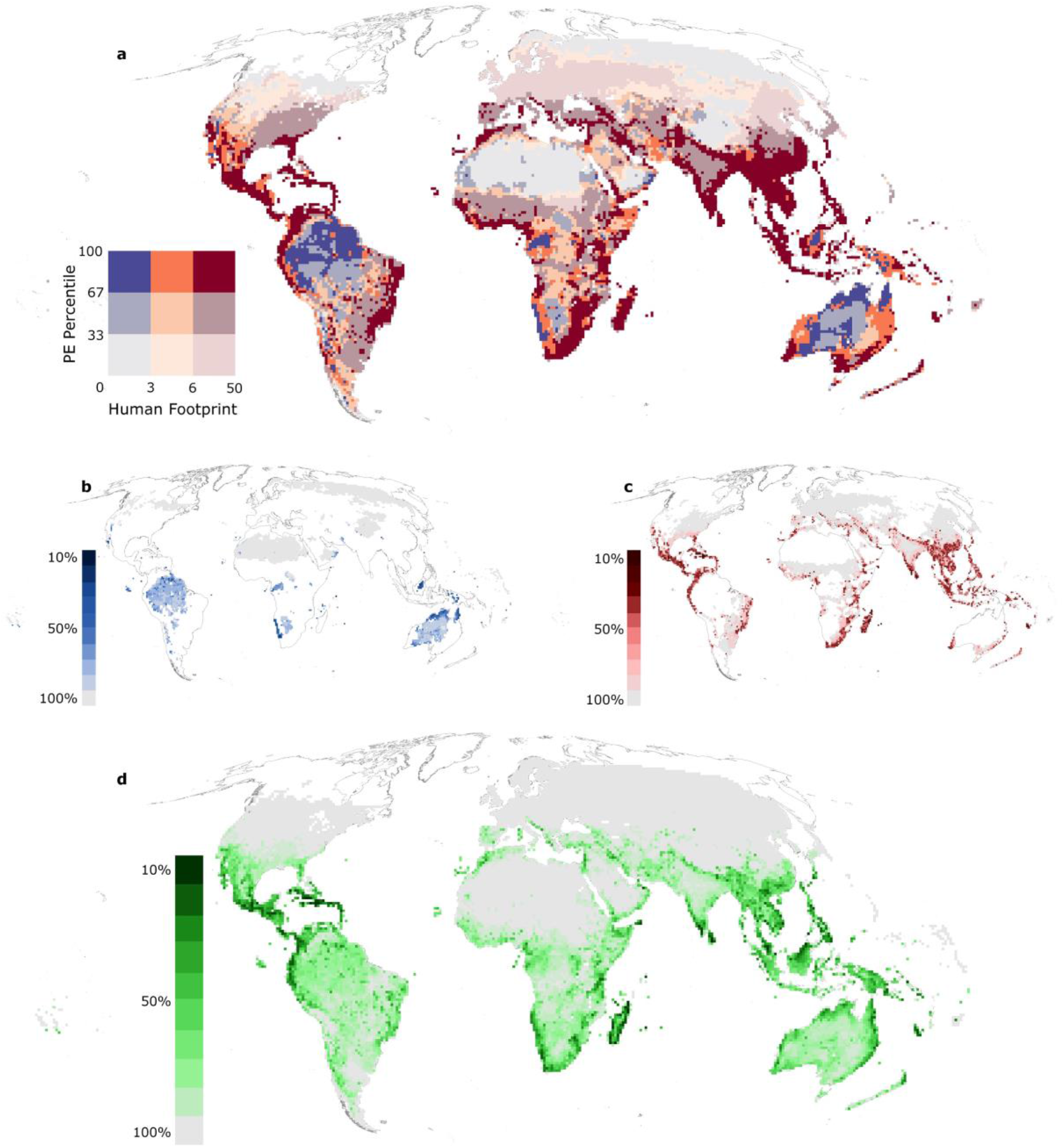
Reptilian Phylogenetic Endemism (PE) and Human Footprint (HF). a) The global relationship between reptilian PE and HF. The colour of the grid cell is determined by HF value and the intensity of the colour by PE percentile. In panels b-d) grid cells are coloured by the cumulative amount of global HIPE captured; darkest blue (b), red (c), and green (d), indicate the highest-scoring grid cells which together capture 10% of global HIPE, whereas the lowest-scoring grid cells also capturing 10% of global HIPE are coloured light grey. Panel b) shows HIPE for low Human Footprint (< 3) grid cells. Panel c) shows HIPE for high Human Footprint cells (≥ 6), and panel d) shows overall global patterns of reptilian HIPE.

Human-Impacted Phylogenetic Endemism (HIPE), is correlated with standard PE for reptiles (r = 0.978, e.d.f. = 448, p ≪ 0.0001; Supplementary Figure 4), despite individual grid cell values differing from PE by up to 1,500% (median = 21%). Highest HIPE value regions (grid cells comprising top 10% of global reptilian HIPE scores) which also coincide with high human pressure (HF ≥ 6) span Southeast Asia, Central America and the Caribbean, Madagascar and Sri Lanka (Figure 2b). Conversely, highest HIPE value regions which also coincide with low human pressure (HF < 3) are restricted to the coast of Namibia, northern Australia and the highlands of Borneo (Figure 2c).

Globally, reptilian HIPE is greatest in Madagascar, Central America and the Caribbean, the Western Ghats of India, Sri Lanka, Socotra, peninsular Malaysia and northern Borneo (Figure 2d). Global patterns of lizard HIPE largely reflect those of all reptiles (Supplementary Figure 5), whereas those for snakes emphasise Central Africa and Southeast Asia (Supplementary Figure 5). High levels of turtle HIPE are concentrated in the Amazon Basin, Central America, southern USA, Southeast Asia, New Guinea, and northern Australia (Supplementary Figure 5).

Grid cells have much greater median and maximum HIPE scores for reptiles than for other tetrapod classes (median = 9.1 x 10^-4^ MY/km^2^ vs amphibians = 4.2 x 10^-4^ MY/km^2^, birds = 4.3 x 10^-4^ MY/km^2^, mammals = 3.6 x 10^-4^ MY/km^2^; maximum = 0.33 MY/km^2^ vs amphibians = 0.30 MY/km^2^, birds = 0.05 MY/km^2^, mammals = 0.03 MY/km^2^). Reptiles contribute a median of 31.1% to tetrapod HIPE scores across all grid cells in which they are present, more than any other class (amphibians = 16.6%, birds =29.7%, mammals = 18%; Supplementary Figure 6). The greatest reptilian contributions (>90% of tetrapod HIPE) occur across the Middle East and North Africa (Figure 3a). The lowest non-zero contributions of reptiles (<10%) occur across northern North America and Europe, the Andes and the Himalayas, where reptiles are scarce.

**Figure 3:**
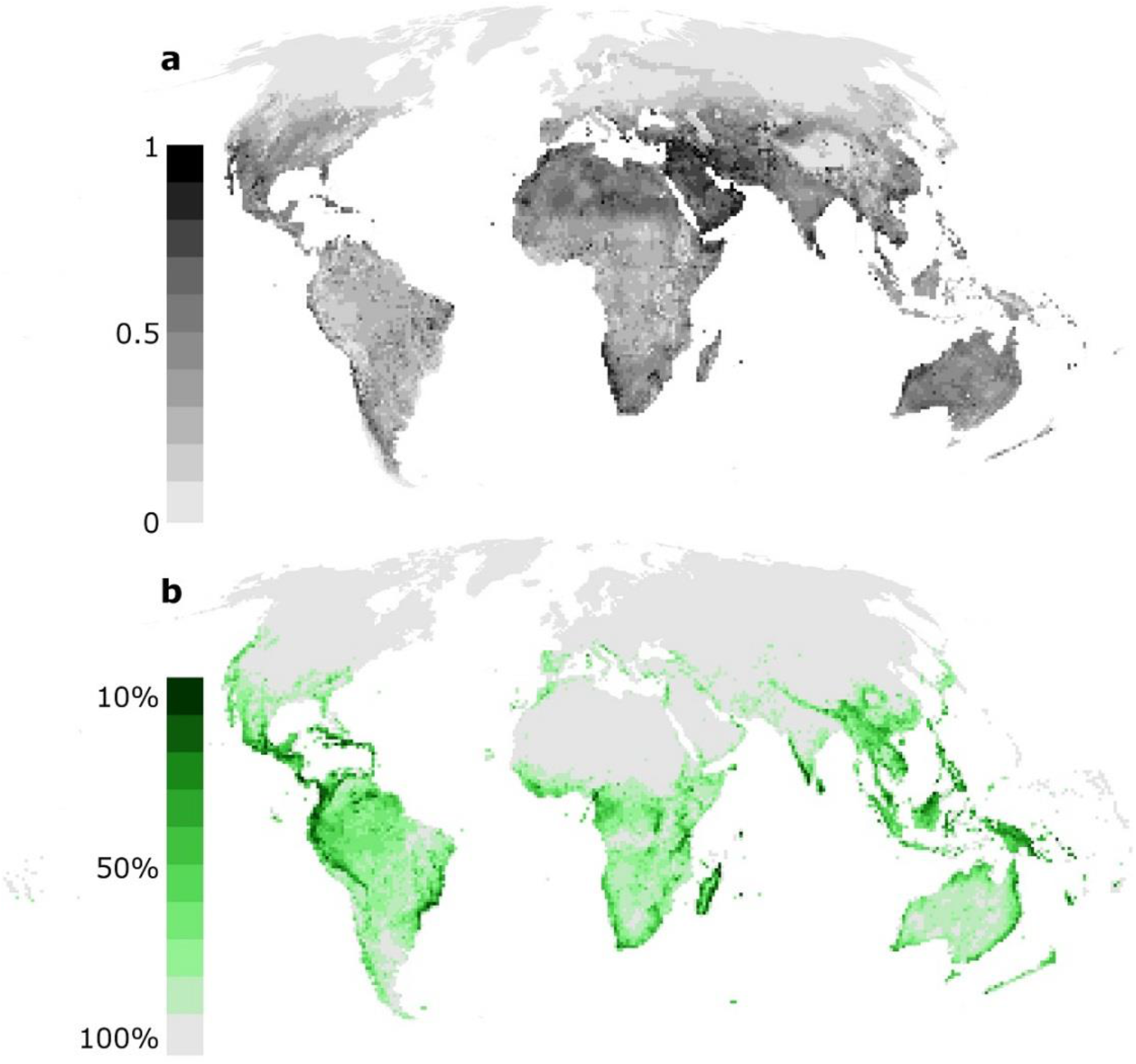
Global patterns of tetrapod HIPE and reptilian contributions. The global patterns of a) the proportion of tetrapod HIPE contributed by reptiles, from 1 (100% of HIPE contributed by reptiles; black) to 0 (0% of HIPE contributed by reptiles; light grey); and b) tetrapod HIPE, where grid cells are coloured by the cumulative amount of global HIPE captured; darkest green cells comprise the highest-scoring grid cells which together capture 10% of global HIPE, whereas the lowest-scoring grid cells which together capture 10% of global HIPE are coloured light grey.

Reptilian HIPE is only moderately correlated with HIPE patterns for other tetrapod classes in each cell across the globe, and inter-correlations are moderate between all classes (Supplementary Figure 7). Turtle HIPE is consistently weakly correlated with that of other reptilian orders and tetrapod classes (r < 0.25, Supplementary Figure 7). Global patterns of tetrapod HIPE are broadly congruent with those for reptiles, but place further emphasis on the importance of the Atlantic coast of Brazil, the Caribbean, Central Africa and New Guinea (Figure 3b). The variation in patterns of clade-specific contributions to global tetrapod HIPE (Figure 3a, Supplementary Figure 6) further highlights the importance of including all tetrapod classes in analyses designed to represent the entire clade.

### Species prioritisation metric for conserving PD

Globally, the 91% of reptiles with phylogenetic data comprise approximately 137 billion years of phylogenetic diversity (PD), significantly more than any other tetrapod class (adjusted p-values from Tukey Honest Significant Differences < 0.0001; amphibians = 130 BY (93% of species), birds = 85 BY (74% of species), mammals = 47 BY (83.5% of species); Supplementary Table 2).

Turtles have the greatest median terminal branch length (TBL) of any tetrapod clade (14.1 MY), whereas lepidosaurs have the greatest maximum TBL (238.7 MY – *Sphenodon punctatus*). Under our new species-level metric, Human-Impacted Terminal Endemism (HITE), lepidosaurs have the second-highest median score (9.9 x 10^-5^ MY/km^2^; lizards = 1.7 x 10^-4^ MY/km^2^, snakes = 2.8 x 10^-5^ MY/km^2^; Figure 4a). This is greater than both birds (3.0 x 10^-5^ MY/km^2^) and mammals (8.5 x 10^-5^ MY/km^2^), with only amphibians scoring higher (median = 5.1 x 10^-4^ MY/km^2^, maximum = 6.3 x 10^-2^ MY/km^2^; Figure 4a).

**Figure 4:**
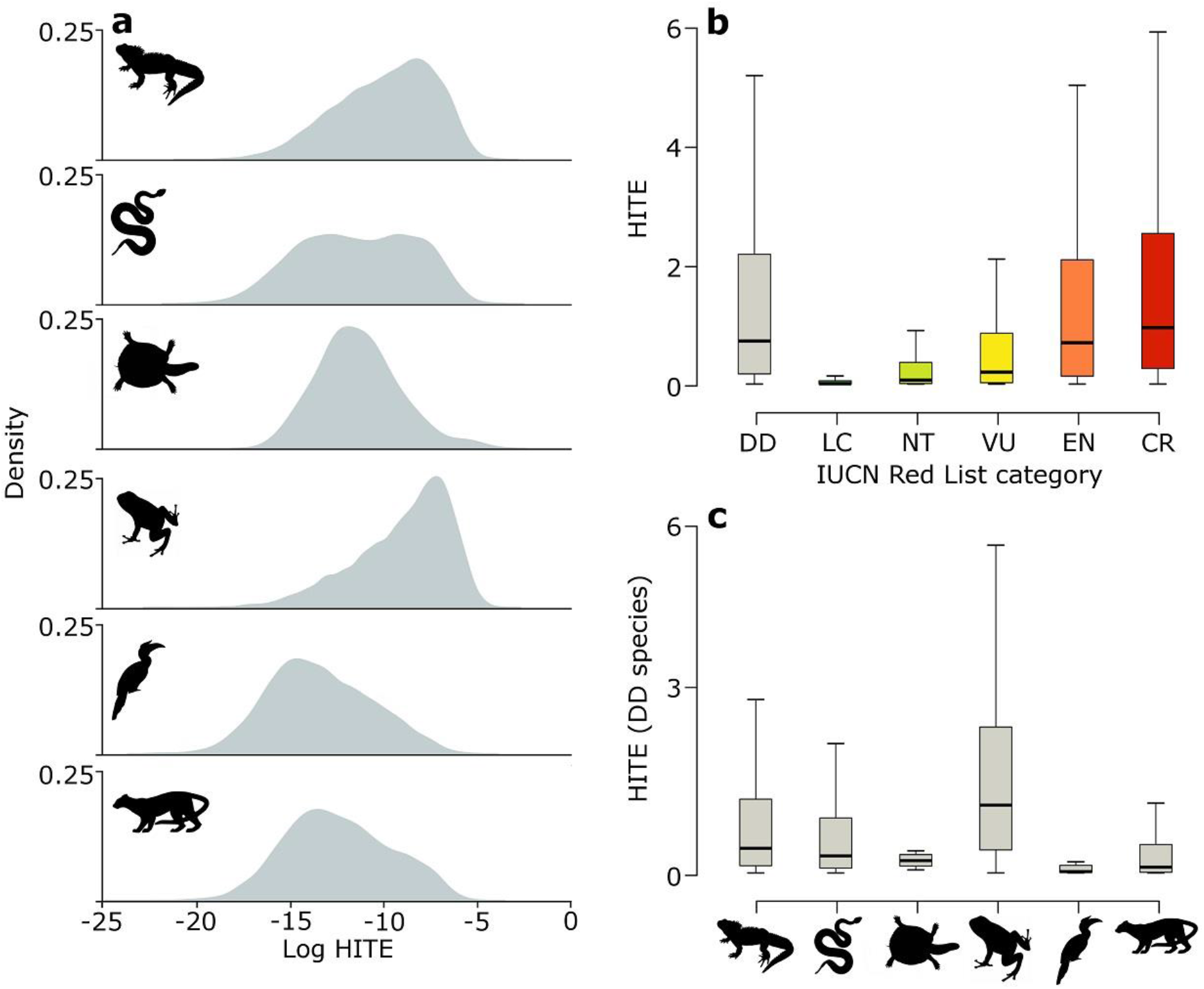
Distributions of Human Impacted Terminal Endemism (HITE) for tetrapods. a) Density distributions of log-transformed HITE scores for tetrapods. Species with long terminal branches occurring in very few grid cells under high human pressure score highly and fall on the right of the x-axis, whereas species with short terminal branches and large ranges encompassing regions of low human pressure fall on the left of the x-axis. Y-axis indicates density of species in each clade with a given HITE value. b) Distribution of HITE scores (in 10^-3^ MY/km^2^) across tetrapods for each IUCN Red List category (excluding Extinct, Extinct in the Wild and unassessed species). c) Distribution of HITE scores (in 10^-3^ MY/km^2^) for Data Deficient (DD) tetrapod species for key tetrapod groups.

Data Deficient tetrapods tend to have longer terminal branches (median = 5.4 MY) than those listed as Least Concern (4.3 MY), Near Threatened (4.3 MY) and Vulnerable (4.8 MY; adjusted p-values from Tukey HSD test < 0.001) and not significantly different from Endangered (5.2 MY) and Critically Endangered (5.5 MY) species (adjusted p-values > 0.05). HITE scores are greater for Data Deficient tetrapods (median = 7.2 x 10^-4^ MY/km^2^) than those listed as Least Concern (6.3 x 10^-6^ MY/km^2^), Near Threatened (6.7 x 10^-5^ MY/km^2^) and Vulnerable (2.0 x 10^-4^ MY/km^2^; adjusted p-values < 0.001), and are comparable to those of Endangered (6.9 x 10^-4^ MY/km^2^) and Critically Endangered species (9.5 x 10^-4^ MY/km^2^; adjusted p-values > 0.05; Figure 4b).

Within Data Deficient species, amphibians have the highest HITE scores (median = 1.5 x 10^-3^ MY/km^2^), followed by lepidosaurs (4.7 x 10^-4^ MY/km^2^; lizards = 5.5 x 10^-4^ MY/km^2^, snakes = 3.8 x 10^-4^ MY/km^2^; Figure 4c). Worryingly, four of the ten highest ranking lizards and eight of the top ten snakes are listed as Data Deficient by the IUCN Red List (ten highest-ranking HITE species for each clade: Supplementary Table 3).

If all reptiles currently listed as threatened by the IUCN Red List were to become extinct (1,196 spp. with phylogenetic data), we stand to lose more than 13.1 billion years of PD (mean; range = 12.3 – 14.3), or around 10% of total reptile PD. This is 1.36 billion years more PD than if extinction risk was randomly distributed across the reptilian phylogeny (paired t-test; t = 20.32, d.f. = 99, p < 0.0001). Given the large proportion of Data Deficient and unassessed reptiles (^~^10% and ^~^34% of all species, respectively), and their potentially high extinction risk, such loss of PD may be much greater, especially where data deficiency for both extinction risk and phylogenetic relationships intersect.

For example, the lizard genus *Dibamus* is represented by 22 species in our study (of the 24 species recognised globally), 16 of which are either unassessed or listed as Data Deficient by the IUCN Red List (as of December 2018). Fifteen of these 22 species are included in the phylogeny despite having no genetic data available, and 12 are known only from their type locality^54^. The amount of PD represented by the 15 species without genetic data is highly uncertain, and ranges from 260 - 1,340 MY across 100 phylogenies (median = 560 MY). Accordingly, estimates of the amount of PD loss due to extinction of unassessed or Data Deficient species range from 0.1 MY (a single species lost with the shortest terminal branch length across 100 phylogenies – 0.00001% additional PD loss) to 1,010 MY (all 16 unassessed/Data Deficient species lost with maximum branch lengths from 100 phylogenies – 7.8% additional PD loss), with a median loss of 230 MY (1.8% additional PD loss).

## Discussion

Globally, reptiles comprise significantly more phylogenetic diversity (PD) than any other tetrapod class. The distribution of reptilian PD largely reflects global richness patterns^21^, though our analysis suggests that extremely high richness in snakes and lizards is achieved through shallow diversification within clades (Figure 1). Our results highlight a large overlap between regions of high human impact and irreplaceable reptilian PD, which is much greater than expected if the two were independent. We therefore incorporated Human Footprint data into our spatial and species-level analyses to capture its potential impact on globally significant concentrations of range-restricted PD. Our metrics represent the first integration of data on environmental pressure affecting terrestrial vertebrates into global prioritisations of imperilled PD.

Reptiles have the highest scores of our spatial metric, Human Impacted Phylogenetic Endemism (HIPE), meaning they are faring worse than amphibians, birds and mammals, and contribute the highest levels of imperilled PD per grid cell. Reptilian contributions to global patterns of tetrapod HIPE are greatest in arid and semi-arid regions, particularly in the Middle East and Southern, North and the Horn of Africa (Figure 2d, Figure 3a)—areas often overlooked in global prioritisations of terrestrial conservation importance for other tetrapod classes^7, 13, 14, 22, 25^. Thus, the inclusion of reptiles in global analyses of this kind is crucial to improve accuracy when attempting to value terrestrial vertebrate diversity for conservation at national, regional and global scales.

Global patterns of tetrapod HIPE emphasise the importance of regions where large amounts of PD are wholly restricted to areas under high human impact; particularly Central America and the Caribbean, Madagascar, the Western Ghats and large swathes of Southeast Asia (Figure 2c), and echo general patterns of Biodiversity Hotspots^55^. These grid cells represent areas of high urgency for conservation of global PD. As HIPE alters the effective range of each species under the assumption that a greater proportion of the range persists in grid cells under lower human impact, it also increases the relative importance of these grid cells. We therefore also highlight areas of the Amazon Basin, the Namib coast of Africa, Central Africa, Northern Australia—regions not captured by existing Biodiversity Hotspots—and the highlands of Borneo (Figure 2b) as long-term conservation priorities, where activities to limit future human impact are more pertinent.

At the species level, reptiles embody more unique evolutionary history than amphibians, birds or mammals. Turtles tend to have particularly long terminal branches, indicating that each turtle species tends to represent large amounts of unique evolutionary history. It is troubling to note that, across tetrapods, Data Deficient and threatened species also generally comprise more unique evolutionary history than non-threatened species. Our species-level metric, Human Impacted Terminal Endemism (HITE), prioritises species with long terminal branches restricted to small ranges under high human impact. Large numbers of small-ranged amphibians and lizards tend to be on long terminal branches and occur in areas of high human impact, and our metric highlights these groups as of major conservation concern.

Many of the highest-ranking HITE tetrapods which have also been classified by the IUCN Red List as Endangered or Critically Endangered are also recognised as priority Evolutionarily Distinct and Globally Endangered (EDGE) species^56^. However, as HITE does not consider IUCN Red List extinction risk data, and uses only phylogeny, range size and Human Footprint, we also identify species of conservation importance which are currently unassessed or listed as Data Deficient by the IUCN. Indeed, we found that Data Deficient tetrapods tend to have HITE scores comparable to those of species listed as Endangered or Critically Endangered. This pattern is particularly pronounced in lizards, snakes and amphibians, where considerably greater proportions of the highest-ranking HITE species for these groups are Data Deficient than either birds or mammals. This suggests that many of the poorly-known amphibians and reptiles are likely to be highly evolutionarily distinct and restricted to regions of intense human pressure. Although such prevalence of high-ranking Data Deficient HITE species is likely driven by higher proportions of data deficiency in amphibians (22%) and reptiles (15%)compared with birds (0.5%) and mammals (14%)^27^, it also highlights the urgent need to assess the extinction risk facing these species in areas of high human impact.

Our case study of the poorly-known lizard genus *Dibamus* underlines the amount of uncertainty we currently face when identifying conservation priorities and estimating impacts of species loss across the tree of life. Our estimation of potential loss of reptilian phylogenetic diversity in this clade varies across four orders of magnitude depending on our assumptions of uncertainty in both phylogeny and extinction risk. Although this is an extreme example, our lack of knowledge of extinction risk and phylogenetic relationships across the reptilian tree of life mean any estimations of potential loss of diversity may be significant underestimates.

It is likely that, without conservation action, we will face losses of billions of years of unique amphibian and reptilian evolutionary history worldwide. While greater research efforts are needed to elucidate the phylogenetic relationships, distribution and population status of poorly known reptiles and amphibians, current and future conservation efforts also need to focus on regions, lineages and species that hold or represent disproportionate amounts of imperilled PD.

## Acknowledgements

J.R. was funded by an Individual Research Fellowship from the Natural Environment Research Council (NERC) (NE/L011611/1). This study is a contribution to Imperial College’s Grand Challenges in Ecosystems and the Environment initiative. R.G. was funded by the Natural Environment Research Council Science and Solutions for a Changing Planet Doctoral Training Programme (grant number NE/L002515/1), the CASE component of which is funded by the Zoological Society of London.

## Author Contributions

R.Gumbs, C.L.G., J.R., N.R.O. conceived the study. R.Gumbs, C.L.G., J.R., M.H., S.M., U.R., W.J. designed the analyses. R.Gumbs conducted the analyses. R.Grenyer, M.B., S.M., U.R. provided reptile spatial data. C.L.G., J.R., M.B., M.H., N.R.O, S.M., U.R., W.J. provided technical support and conceptual advice. C.L.G., J.R., N.R.O. supervised the study. R.Gumbs wrote the paper, with substantial contributions from J.R, C.L.G., M.B., M.H, N.R.O., S.M., U.R., W.J.

