## Supplementary material for "Global priorities for conservation of reptilian phylogenetic diversity in the face of human impacts"

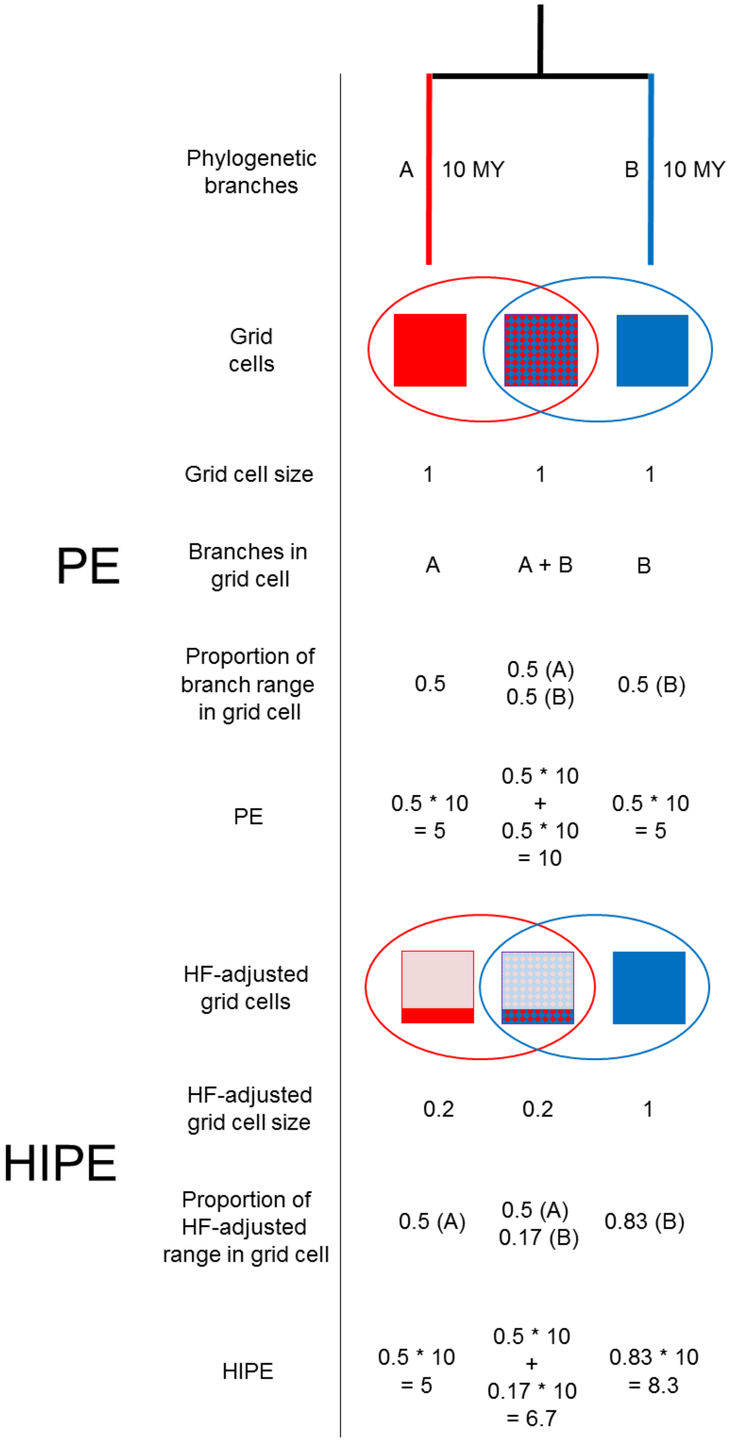


**Supplementary Figure 1: HIPE and PE calculated for a simple example.** The HIPE and PE scores for three grid cells. Each grid cell is coloured according to which branch is present: blue = branch A, red =

branch B, and blue + red = branches A and B. The proportion of saturated vs. faded colour corresponds to the Human Footprint-adjusted grid cell size of each cell (smaller HF-adjusted range size = higher human impact). The coloured circles encompass the grid cells in which each branch is present. Under PE, both branch A and B are present in two grid cells, thus each of the three grid cells comprises 0.5 of all cells occupied by the branch. This changes when Human Footprint is incorporated due adjust effective range size, and HIPE scores differ from those of PE according to this shift in proportion of effective range size represented by a grid cell.

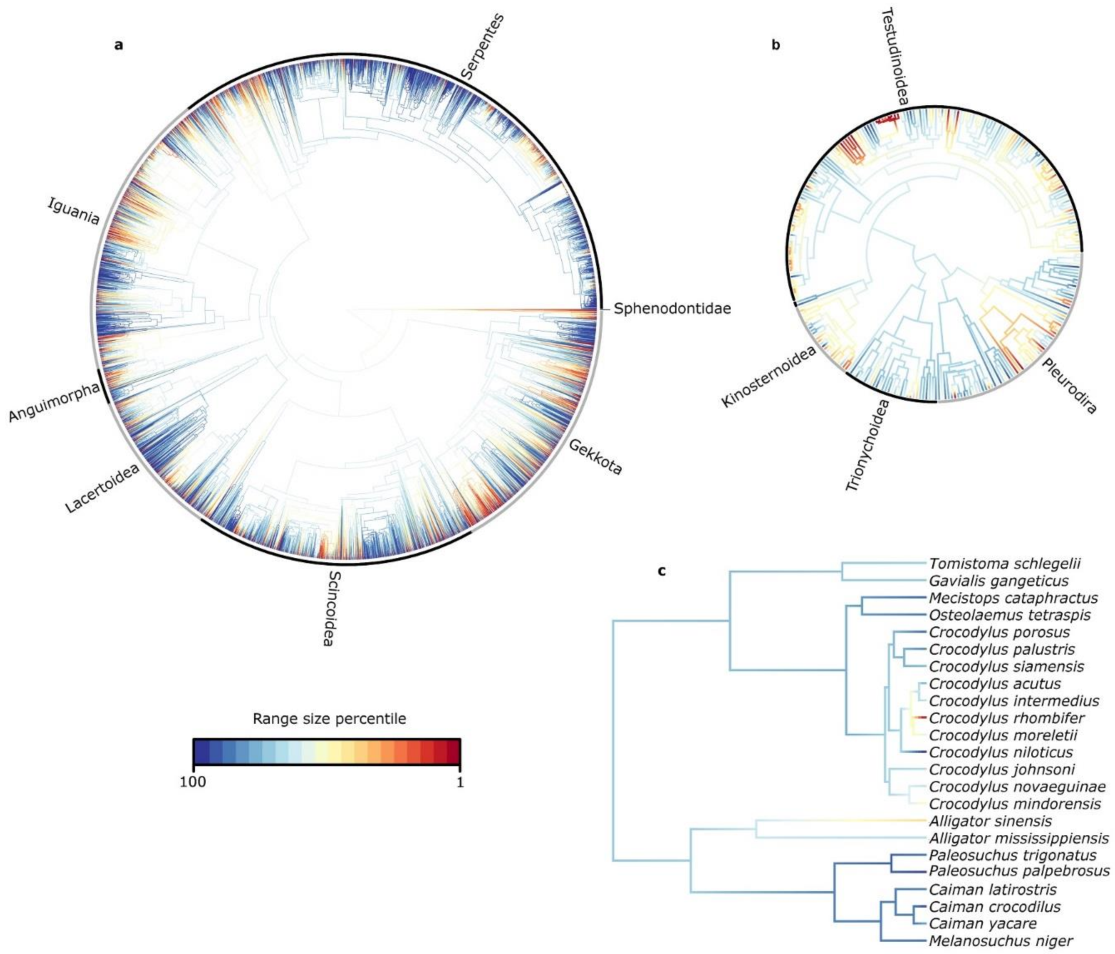


**Supplementary Figure 2: Phylogenetic distribution of range size across reptiles.** The percentiles of range size, measured in number of grid cells, across the phylogenies of a) lepidosaurs (squamates + tuatara), b) testudines, and c) crocodilians.

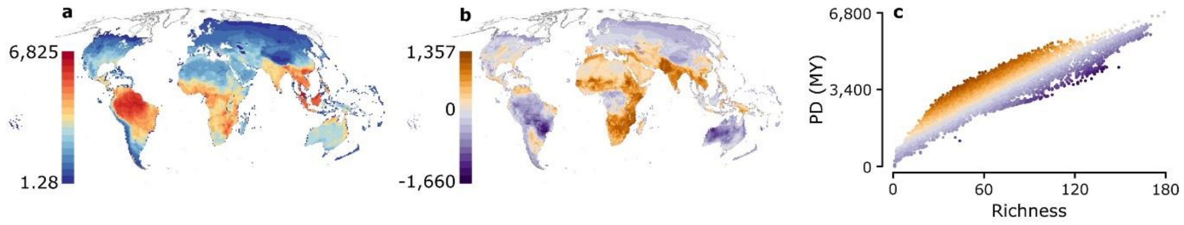


Supplementary Figure 3: Global patterns of reptilian phylogenetic diversity (PD). Cumulative PD (a), amount of PD per grid cell greater or lower than expected for the observed species richness (b), and the relationship between richness and PD across all grid cells (c) for all reptiles.

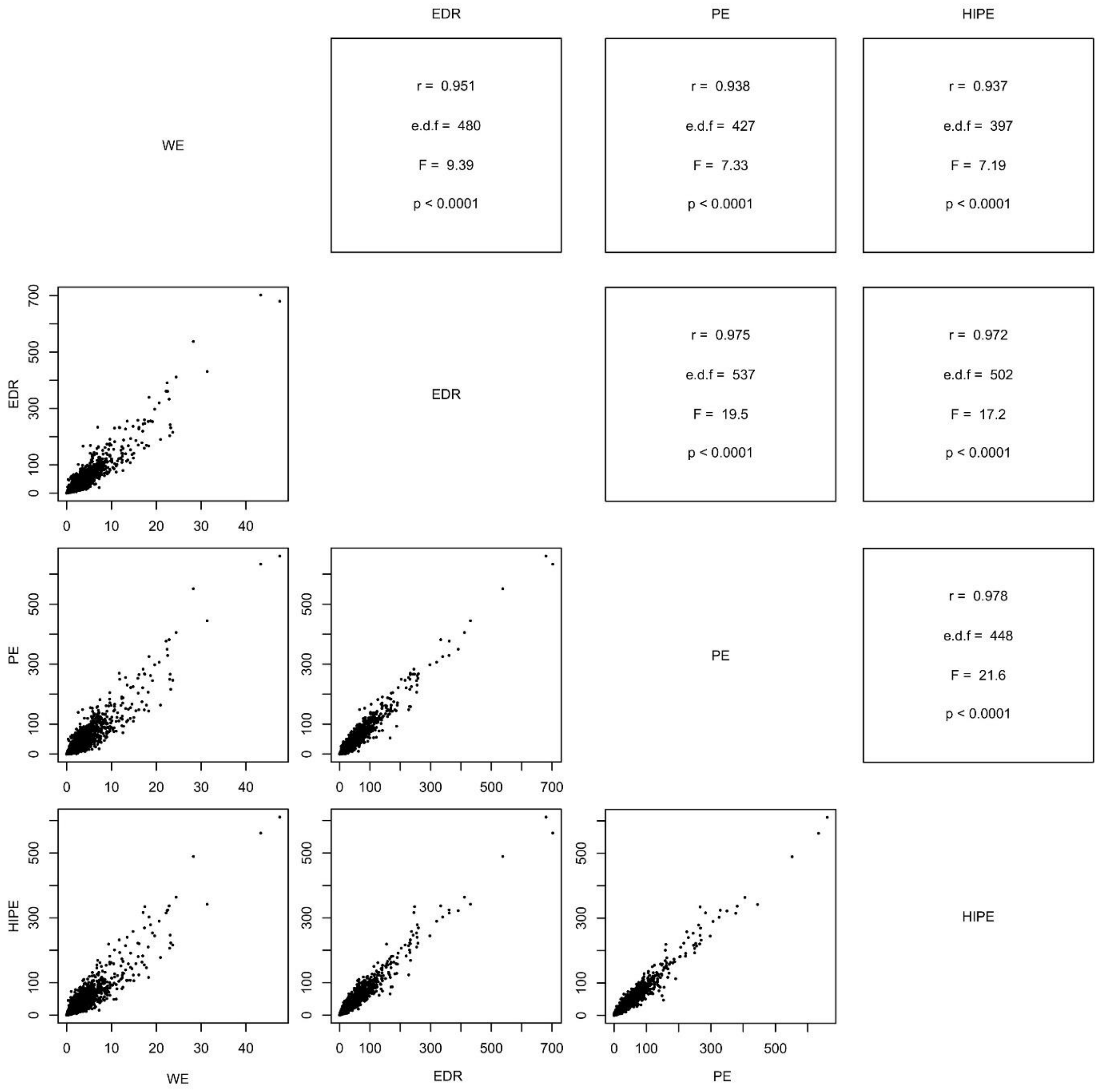


**Supplementary Figure 4: Relationships of spatial metrics for all reptiles.** Results of spatially corrected Pearson's correlations among WE, EDR, PE and HIPE (above diagonal split) and scatterplots of the values for each grid cell of global reptile distribution for each metric (below diagonal split).

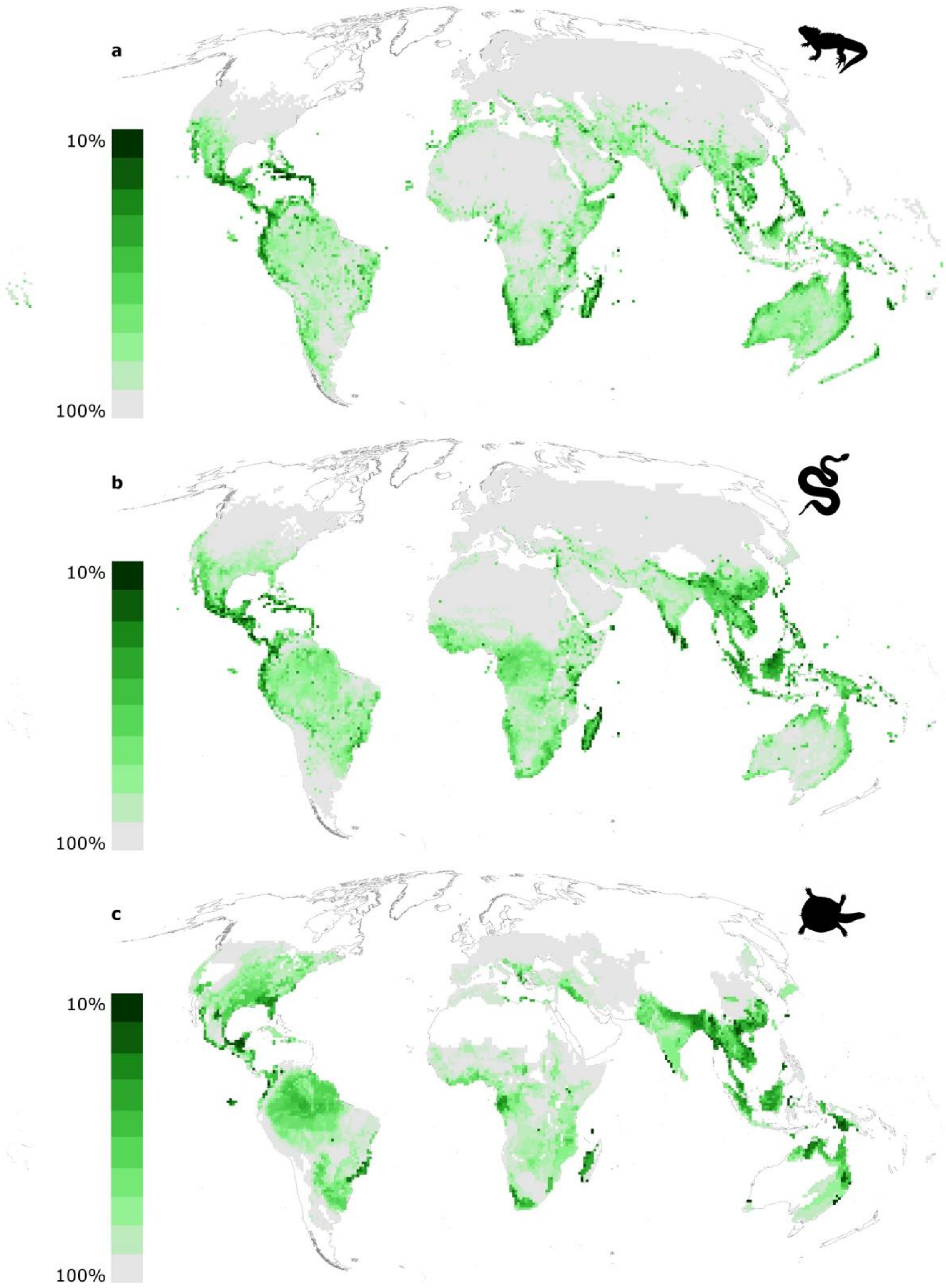


**Supplementary Figure 5: Global patterns of HIPE for reptilian clades.** The global patterns of HIPE for a) lizards (lizards, amphisbaenians and the tuatara), b) snakes and c) testudines. The top 10% ranked grid cells for HIPE are darkest green and the lowest ranking 10% are coloured light grey.

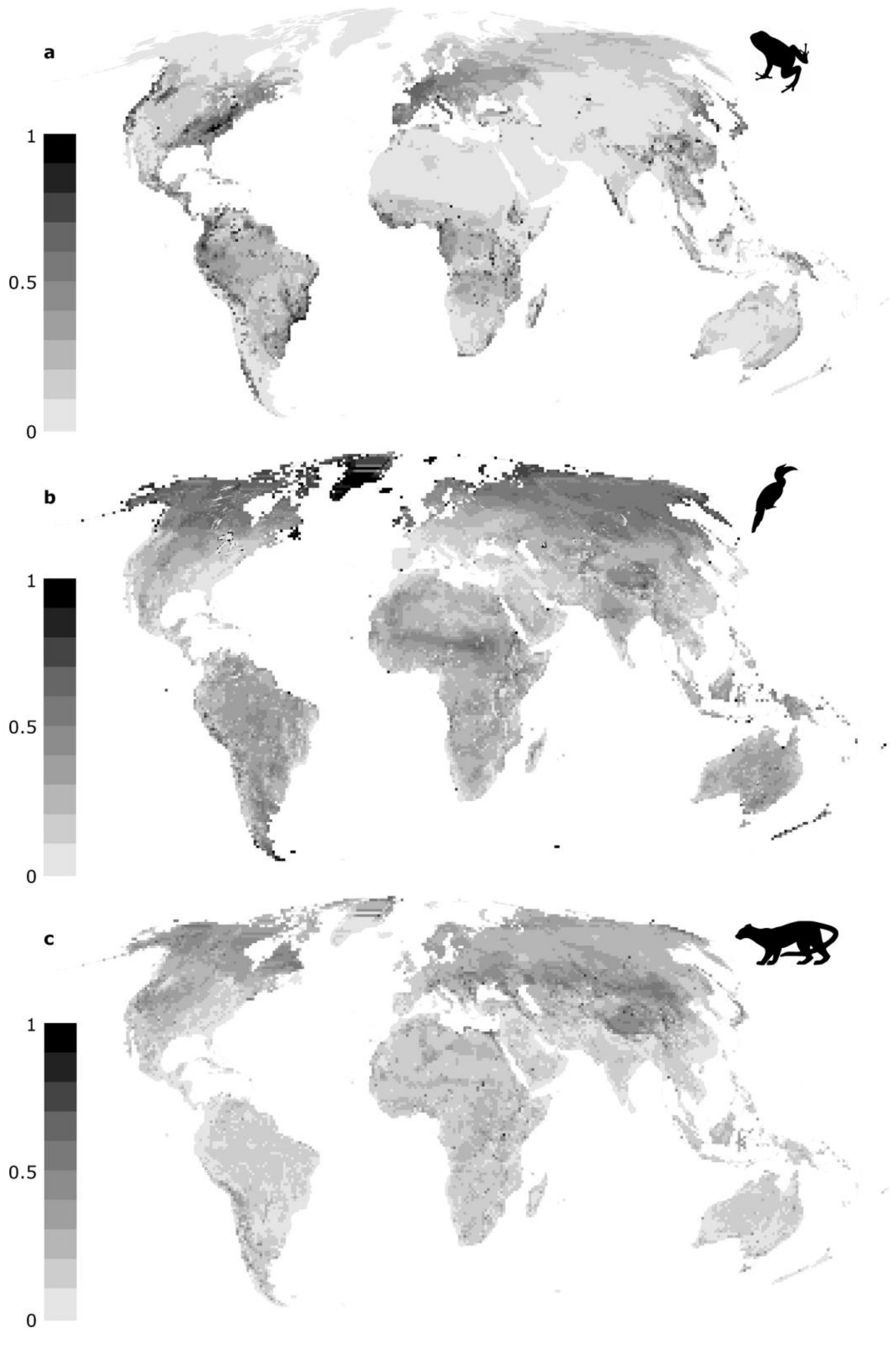


Supplementary Figure 6: Global contributions to tetrapod HIPE by non-reptilian tetrapods. The proportional contributions to tetrapod HIPE scores by a) amphibians, b) birds and c) mammals. Grid cell scores range from 1 (100% of HIPE contributed by clade; black) to 0 (0% of HIPE contributed by clade; light grey).

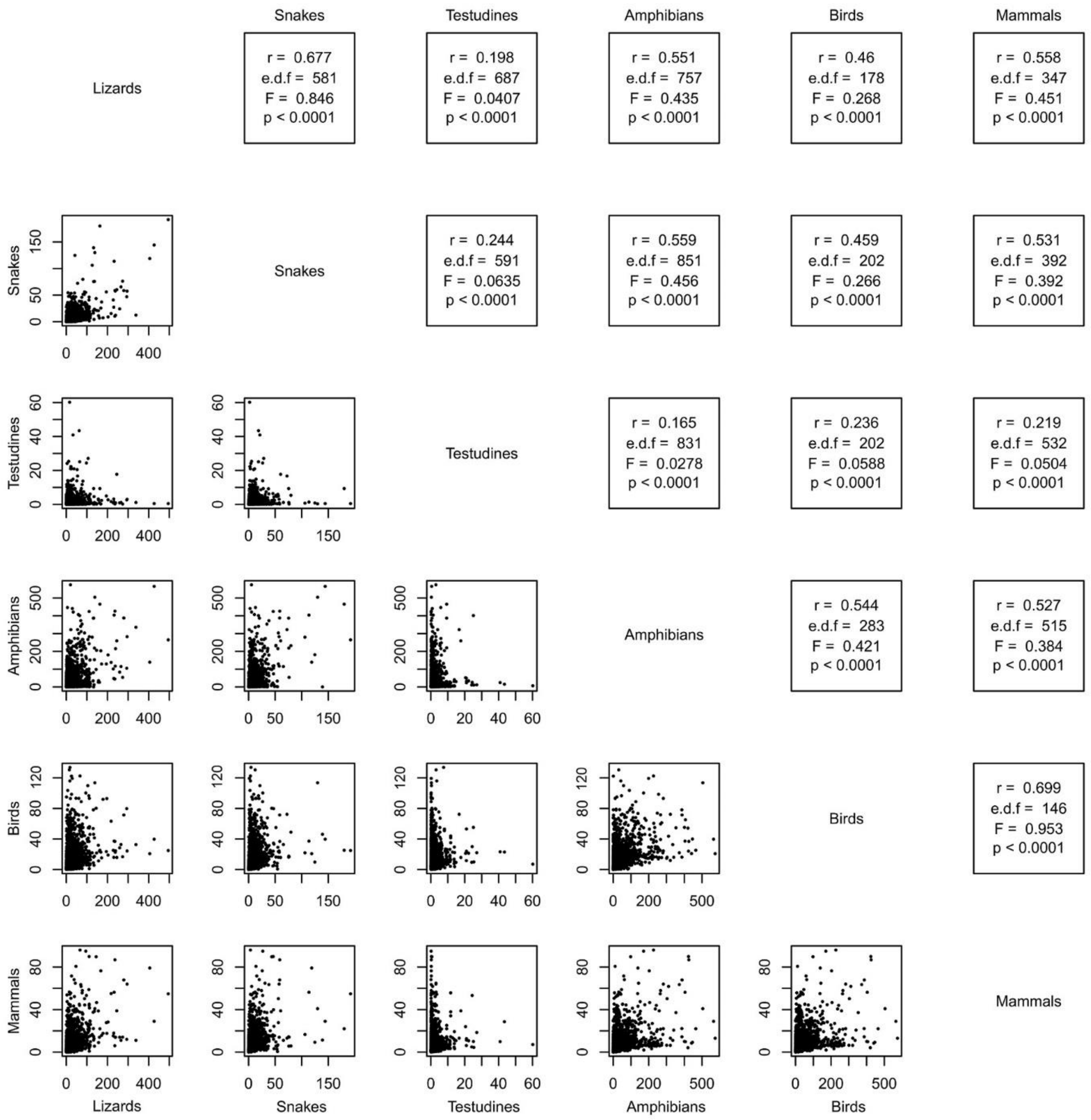


**Supplementary Figure 7: relationships amongst reptile and tetrapod groups for all grid cells of global HIPE.** Results of spatially corrected Pearson's correlations for between HIPE scores of tetrapod groups (above diagonal split) and scatterplots of the values for each non-zero grid cell of global HIPE values for each tetrapod group (below diagonal split).

**Supplementary Table 1: Taxonomic representation for each reptilian order and tetrapod class in this study.** Number and percentage of species for each clade with both spatial and phylogenetic data available. Data sources in Supplementary References.

| Clade | Total PD (MY) (% of described species in phylogeny)* | Species with phylogenetic and range data | Percentage of total species (total number of species) |
|---|---|---|---|
| Reptiles | 136,962 (91%) | 9,862 | 90.9% (10,845)[1] |
| Crocodilians | 531 (95.8%) | 23 | 95.8% (24)[1] |
| Testudines | 8,213 (80.4%) | 282 | 80.3% (351)[1] |
| Lepidosaurs | 128,218 (91.3%) | 9,557 | 91.3% (10,470)[1] |
| Amphibians | 130,703 (93.1%) | 5,874 | 75.5% (7,776)[2] |
| Birds | 85,469 (91.1%) | 9,274 | 84.5% (10,970)[3] |
| Mammals | 46,649 (83.5%) | 4,386 | 77% (~5,692)[4] |
| **Tetrapods** | **399,783 (91.1%)** | **29,396** | **84.2% (~34,906)** |

*Median value taken from random sample of 100 phylogenies for all clades except testudines and crocodilians, for which only single consensus phylogenies were available

**Supplementary Table 2. Spatial metrics comparison.** A simple description, calculation, and worked example for each of the three base metrics with which we compare our novel spatial metric.

| Metric | Description | Calculation | Example |
|---|---|---|---|
| Weighted Endemism (WE)[5,6] | WE is a metric that applies to a grid cell. It captures how critical the cell is for its contributions to the total range size of the species it | $WE_i$ , the weighted endemism of cell $i$ is given by $$WE_i = \sum_{j=1}^{S} q_{ij}$$ | For a focal grid cell containing two species, A and B. Species A occurs in a total of 5 grid cells globally, including our focal cell. |

| | | | |
|---|---|---|---|
| | contains. WE is calculated as a sum over all species in the grid cell. Each species' contribution to WE is given by the fraction of its total range that lies within the cell. | Where $q_{ij}$ is the fraction of the range (in grid cells) of species *j* found in grid cell *i* and S is the number of species. The sum can be considered as a sum over all species as $q_{ij} = 0$ if species *j* is not found in grid cell *i.* | Species B occurs in a total of 4 grid cells globally, including our focal cell. Species A contributes (1/5) to WE and species B contributes (1/4) . This gives WE = (1/5) + (1/4) = 0.45 (units are grid cell$^{-1}$). |
| Evolutionary Distinctness Rarity (EDR)[7] | EDR is a metric for a given species. Specifically, it measures unique evolutionary history (Evolutionary Distinctiveness - ED) of the species weighted by the species' rarity as inferred by its total range size. | $EDR_j$, the EDR score of a species *j* is given by $$EDR_j = ED_j \cdot q_{ij}$$ Where $ED_j$ is the Evolutionary Distinctiveness of species *j* and $q_{ij}$ is the fraction of the distribution (in grid cells) of species *j* found in cell *i*, which should be one of its occupied cells, in practice $q_{ij}$ is given by the reciprocal of the species' range size ($1/R_j$). | For a species that occurs in 5 grid cells and has an ED of 10 MY (million years), that species has an EDR of 10/5 = 2 (units are MY/grid cell). |
| Phylogenetic Endemism (PE)[5] | An extension of Weighted Endemism which distributes the length of each phylogenetic branch equally across all grid cells in which that phylogenetic branch is found. Measures the amount of phylogenetic diversity (PD) represented by each grid cell, assuming | $PE_i$, the Phylogenetic Endemism score of cell *i* is given by $$PE_i = \sum_{b=1}^{2S-1} L_b q_{i,b}$$ Where $L_b$ is the length of phylogenetic branch *b*, S is the species richness in cell *i* so that *2S -1* is the total number of branches and $q_{i,b}$ is the total proportion | For a grid cell containing three phylogenetic branches: one branch unique to species A (branch A), one branch unique to species B (branch B), both of which are 8 MY in length, and a third branch from which both species are descended (branch C), |

| | | | |
|---|---|---|---|
| | total PD is distributed evenly in space. Global PD is given by the sum of PE across all global grid cells. | of the range of branch *b* that falls within cell *i*. $q_{i,b} = 0$ if the branch has no descendent species in cell *i*. | which is 4 MY in length. Branch A occurs in 5 grid cells, branch B in 4 cells, but both species co-occur in two grid cells, thus branch C occurs only in (5+4-2 = 7) grid cells. Thus PE = (8/5)+(8/4)+(4/7) = 4.17 MY/grid cell |
| Human-Impacted Phylogenetic Endemism (HIPE) | An extension of Phylogenetic Endemism which scales the size of each grid cell based on the level of human impact in that cell to provide a 'Human Footprint-adjusted range size'. Phylogenetic Endemism is then calculated based on these Human Footprint-adjusted range sizes for each branch of the tree rather than on the pure geographic range size. Less impacted cells thus receive a higher score if ranges are spread across a landscape with variable levels of human impact. Where the human impact is equal for all cells in an | $HIPE_i$ the Human-Impacted Phylogenetic Endemism of cell *i* is given by $HIPE_i = \sum_{b=1}^{2S-1} L_b \times \frac{H_{i,j}}{HR_j}$ Where $L_b$ is the length of each phylogenetic branch *b*, $H_{i,j}$ is the Human Footprint-adjusted range of species *j* in *i* (zero if the species is absent from cell i) and $HR_j$ is the total Human Footprint-adjusted range size of species *j* across all cells. | For a grid cell with a HF-adjusted range size of 0.8, containing three phylogenetic branches: one branch unique to species A (branch A), one branch unique to species B (branch B), both of which are 8 MY in length, and a third branch from which both species are descended (branch C), which is 4 MY in length. Branch A occurs in 5 grid cells, each with a HF-adjusted range size of 0.8. Branch B occurs in 4 cells, two which it shares with Branch A (HF-adjusted range size = 0.8) and two with a HF-adjusted range size of 0.2. Branch C therefore occurs only in |

| | | | |
|---|---|---|---|
| | analysis, HIPE gives the same result as PE. | | seven grid cells, 5 of which have a HF-adjusted range size of 0.8 and two of 0.2. HIPE = (8*(0.8/(5*0.8))) + (8*(0.8/(2*0.2+2*0.8))) + (4*(0.8/(5*0.8+2*0.2))) = 5.527 MY/grid cell |
| Human Impacted Terminal Endemism (HITE) | A species level measure chosen to incorporate the common elements of EDR and PE. HITE represents the terminal branch length of a taxa scaled by a measure of its rarity, given by the reciprocal of its Human Footprint-adjusted range size. | $HITE_j$, the HITE score of a species *j* is given by $$HITE_j = TBL_j \cdot \frac{1}{HR_j}$$ Where $TBL_j$ is the Terminal Branch Length of species *j* in the phylogeny, and $HR_j$ is the total Human Footprint-adjusted range size of species *j* across all grid cells. | For a species that occurs in 5 grid cells (two cells with an HF-adjusted size of 0.2 and three cells with an HF-adjusted size of 0.6), then the species has a total Human Footprint-adjusted range size of (2*0.2 + 3 *0.6). Suppose the species has a terminal branch length of 7 MY (million years), that species has an HITE score of 7/(2*0.2 + 3 *0.6) = 3.182 (units are MY/grid cell). |

**Supplementary Table 3: The ten highest ranking HITE species for each tetrapod group.** The ten species with the largest Human Impacted Terminal Endemism (HITE) scores for each group and their IUCN Red List status as of December 2018. NE = Not Evaluated, DD = Data Deficient, LC = Least Concern, NT = Near Threatened, VU = Vulnerable, EN = Endangered, CR = Critically Endangered.

| Species | HF-adjusted range size | TBL | HITE | IUCN Red List Status |
|---|---|---|---|---|
| Lizards | | | | |

| | | | | |
|---|---|---|---|---|
| *Dibamus somsaki* | 0.4 | 140.2 | 350.4 | DD |
| *Dibamus dalaiensis* | 0.4 | 119.1 | 297.7 | LC |
| *Goniurosaurus kuroiwae* | 0.2 | 53.7 | 268.5 | VU |
| *Gekko canaensis* | 0.2 | 52.1 | 260.7 | LC |
| *Brachymeles wrighti* | 0.2 | 50.2 | 251.1 | DD |
| *Cricosaura typica* | 0.4 | 74.4 | 186.1 | NT |
| *Luperosaurus yasumai* | 0.2 | 36.8 | 184.1 | DD |
| *Dibamus vorisi* | 0.4 | 71.2 | 178.0 | DD |
| *Cnemaspis psychedelica* | 0.2 | 33.0 | 165.2 | EN |
| *Gonatodes daudini* | 0.2 | 33.0 | 165.0 | CR |
| Snakes | | | | |
| *Gerrhopilus bisubocularis* | 0.2 | 49.2 | 245.8 | DD |
| *Epictia rubrolineata* | 0.2 | 26.7 | 133.5 | DD |
| *Gerrhopilus oligolepis* | 0.2 | 25.0 | 125.2 | DD |
| *Bitia hydroides* | 0.2 | 23.3 | 116.3 | LC |
| *Tricheilostoma greenwelli* | 0.2 | 22.0 | 109.9 | DD |
| *Gerrhopilus tindalli* | 0.2 | 18.5 | 92.4 | DD |
| *Oligodon travancoricus* | 0.2 | 17.2 | 86.0 | DD |
| *Pareas nigriceps* | 0.4 | 33.5 | 83.7 | DD |
| *Opisthotropis tamdaoensis* | 0.2 | 15.0 | 75.2 | DD |
| *Tetracheilostoma bilineatum* | 0.4 | 30.0 | 75.0 | LC |
| Testudines | | | | |
| *Pseudemydura umbrina* | 0.4 | 89.6 | 223.9 | CR |
| *Geoemyda japonica* | 0.2 | 28.5 | 142.6 | EN |
| *Elusor macrurus* | 0.4 | 37.3 | 93.1 | EN |
| *Astrochelys yniphora* | 0.4 | 30.8 | 76.9 | CR |
| *Siebenrockiella leytensis* | 0.4 | 30.4 | 76.1 | CR |
| *Pyxis planicauda* | 0.4 | 16.5 | 41.1 | CR |
| *Myuchelys georgesi* | 0.6 | 21.7 | 36.2 | DD |
| *Myuchelys purvisi* | 2.6 | 53.8 | 20.7 | NE |
| *Pyxis arachnoides* | 0.8 | 16.5 | 20.6 | CR |
| *Pelusios broadleyi* | 0.6 | 9.1 | 15.2 | VU |
| Amphibians | | | | |
| *Chikila fulleri* | 0.2 | 117.5 | 587.5 | DD |
| *Karsenia koreana* | 0.2 | 84.5 | 422.5 | LC |
| *Nasikabatrachus sahyadrensis* | 0.4 | 145.5 | 363.7 | EN |
| *Phytotriades auratus* | 0.2 | 56.5 | 282.7 | CR |
| *Latonia nigriventer* | 0.2 | 56.5 | 282.3 | CR |
| *Ptychadena filwoha* | 0.4 | 110.1 | 275.4 | DD |
| *Eleutherodactylus counouspeus* | 0.2 | 53.2 | 266.2 | EN |
| *Platymantis isarog* | 0.2 | 52.3 | 261.3 | LC |
| *Micrixalus narainensis* | 0.2 | 51.0 | 255.1 | DD |
| *Scinax muriciensis* | 0.2 | 49.1 | 245.6 | CR |
| Birds | | | | |

| | | | | |
|---|---|---|---|---|
| *Microeca hemixantha* | 0.2 | 23.7 | 118.3 | NT |
| *Nipponia nippon* | 0.2 | 20.7 | 103.6 | EN |
| *Regulus madeirensis* | 0.2 | 17.9 | 89.6 | LC |
| *Zeledonia coronata* | 0.2 | 17.5 | 87.7 | LC |
| *Circus maillardi* | 0.2 | 12.8 | 64.1 | EN |
| *Papasula abbotti* | 0.2 | 12.5 | 62.4 | EN |
| *Nesoenas mayeri* | 0.2 | 12.2 | 61.1 | VU |
| *Nesillas mariae* | 0.2 | 11.8 | 59.2 | LC |
| *Dicaeum quadricolor* | 0.2 | 10.6 | 53.1 | CR |
| *Leucocarbo carunculatus* | 0.2 | 10.5 | 52.5 | VU |
| Mammals | | | | |
| *Calcochloris tytonis* | 0.2 | 27.6 | 138.0 | DD |
| *Myrmecobius fasciatus* | 0.4 | 30.5 | 76.3 | EN |
| *Spalax arenarius* | 0.2 | 15.1 | 75.4 | EN |
| *Thomomys bulbivorus* | 0.2 | 14.5 | 72.5 | LC |
| *Gymnobelideus leadbeateri* | 0.4 | 25.9 | 64.8 | CR |
| *Hipposideros inexpectatus* | 0.4 | 25.7 | 64.3 | DD |
| *Niviventer culturatus* | 0.2 | 12.3 | 61.3 | LC |
| *Crocidura wimmeri* | 0.2 | 11.6 | 57.8 | CR |
| *Mus famulus* | 0.2 | 11.2 | 56.0 | EN |
| *Crocidura orientalis* | 0.2 | 10.9 | 54.5 | LC |
